# Magia: Robust automated image processing and kinetic modeling toolbox for PET neuroinformatics

**DOI:** 10.1101/604835

**Authors:** Tomi Karjalainen, Jouni Tuisku, Severi Santavirta, Tatu Kantonen, Lauri Tuominen, Jussi Hirvonen, Jarmo Hietala, Juha O. Rinne, Lauri Nummenmaa

## Abstract

**Introduction:** Modelling of the radioactivity images produced by PET scanners into biologically meaningful quantities, such as binding potential, is a complex multi-stage process involving data retrieval, preprocessing, drawing reference regions, kinetic modelling, and post-processing of parametric images. The process is challenging to automatize mainly because of manual work related to input generation, thus prohibiting large-scale standardized analysis of brain PET data. To resolve this problem, we introduce the Magia pipeline that enables processing of brain PET data with minimal user intervention. We investigated the accuracy of Magia in the automatic brain-PET data processing with four tracers binding to different binding sites: [_11_C]raclopride, [_11_C]carfentanil, [_11_C]MADAM, and [_11_C]PiB.

**Materials and methods:** For each tracer, we processed 30 historical control subjects’ data with manual and automated methods. Five persons manually delineated the reference regions (cerebellar or occipital cortex depending on tracer) for each subject according to written and visual instructions. The automatic reference-region extraction was based on FreeSurfer parcellations. We first assessed inter-operator variance resulting from manual delineation of reference regions. Then we compared the differences between the manually and automatically produced reference regions and the subsequently obtained metrics.

**Results:** The manually delineated reference regions were remarkably different from each other. The differences translated into differences in outcome measures (binding potential or SUV-ratio), and the intra-class correlation coefficients were between 47 % and 96 % for the tracers. While the Magia-derived reference regions were topographically very different from the manually defined reference regions, Magia produced outcome measures highly consistent with average of the manually obtained estimates. For [_11_C]carfentanil and [_11_C]PiB there was no bias, while for [_11_C]raclopride and [_11_C]MADAM Magia produced 3-5 % higher binding potentials as a result of slightly lower time-integrals of reference region time-activity curves.

**Conclusion:** Even if Magia produces reference regions that are anatomically different from manually drawn reference regions, the resulting outcome measures are highly similar. Based on these results and considering the high inter-operator variance of the manual method, the high level of standardization and strong scalability of Magia, we conclude that Magia can be reliably used to process brain PET data.

## Introduction

Statistical power of neuroimaging studies has been widely questioned in the recent years, leading to calls for significantly larger samples are required to avoid false positive and negative findings ^1–3^. Additionally, the role of researcher degrees of freedom, i.e. the subjective choices made during the process from data collection to its analysis, has been identified as an important reason for poor replicability of many findings. ^4^ Consequently, the focus in neuroimaging has shifted towards standardized, large-scale neuroinformatics based approaches ^5, 6^. Today, several standardized and highly automatized preprocessing pipelines are publicly available for processing functional magnetic resonance images (fMRI) ^7^. Such standardized methods are not, however, currently widely used for analysis of positron emission tomography (PET) data.

Compared to fMRI preprocessing, preprocessing of PET data is relatively straightforward because confounding temporal signals are rarely regressed out of the data, and the preprocessing thus only consists of spatial processes, such as frame-realignment and coregistration. Yet, any all-inclusive PET processing pipeline must be able to handle numerous kinetic models to support as many radiotracers as possible. Thus, unlike fMRI preprocessing tools, PET pipelines should handle both the preprocessing as well as the kinetic modeling. Presumably, this is the reason why such systems have not been available. A particularly sensitive task in PET analysis is the requirement of input function. Depending on tracer, the input function can be obtained either from blood samples or directly from the PET images if a reference region is available for the tracer. The blood samples require manual processing before the input function can be obtained from them. While population-based atlases ^8–10^ provide an automatic way for defining reference regions ^11–13^, they are suboptimal because the process requires spatial normalization of the images. Ideally, the reference region should be defined separately for each individual before spatial normalization. Consequently, manual delineation is still considered the golden standard for defining the reference regions, thus prohibiting fully automatic analysis of PET data. Furthermore, manual reference region delineation is time-consuming and relies on numerous subjective choices. To minimize between-study variance resulting from operator-dependent choices ^14^, a single individual should delineate the reference regions for all studies within a project. Thus, manual delineation is not suited for large-scale projects where hundreds of scans are processed, or neuroinformatics approaches where even significantly larger number of scans have to be processed.

To resolve these problems, we have introduced the Magia analysis pipeline that enables automatic modelling of brain PET data with minimal user intervention (https://github.com/tkkarjal/magia). The major advantages of this approach involve:

1. Flexible, parallelizable environment suitable for large-scale standardized analysis
2. Fully automated processing of brain PET data starting from raw images.
3. Visual quality control of the processing steps.
4. Centralized management and storage of study metadata, image processing methods and outputs for subsequent reanalysis and quality control.

In this study we compared Magia-derived input functions and the subsequent outcome measures against those obtained using conventional manual techniques with four tracers binding to different sites: [_11_C]raclopride, [_11_C]carfentanil, [_11_C]MADAM, and [_11_C]PiB. We also assessed inter-rater agreement in the reference region definition and uptake estimates, and regional and voxel-level outcome measures.

## Materials and methods

### Overview of Magia

Magia (https://github.com/tkkarjal/magia) is a fully automatic analysis pipeline for brain PET data. Running on MATLAB (The MathWorks, Inc., Natick, Massachusetts, United States), Magia combines methods from SPM (www.fil.ion.ucl.ac.uk/spm/) and FreeSurfer (https://surfer.nmr.mgh.harvard.edu/) as well as in-house software developed for kinetic modeling. Magia has been developed alongside a centralized database (http://aivo.utu.fi) containing metadata about each study, facilitating data storage and neuroinformatics-type large-scale PET analyses. While implementation of a similar database is highly recommended, Magia can also be installed and used without such database as long as the user can feed in the necessary information about the studies. Magia runs only on Linux/Mac. The Optimization Toolbox for MATLAB is required for fitting some of the models. Magia has been developed using MATLAB R2016b. Magia currently supports the simplified reference tissue model, Logan with both plasma input and reference tissue input, Patlak with both plasma input and reference tissue input, SUV-ratio, and FUR analysis for late scans with plasma input. Also two-tissue compartmental model can be fitted to ROI-level data.

A box-diagram describing the main steps in Magia processing is shown in Figure 1. Magia starts by preprocessing the PET images. The preprocessing consists of frame-alignment and coregistration with the MRI. The MRI is processed with FreeSurfer to generate anatomical parcellations for defining regions of interest ^12^, including a reference region if one is available for a tracer. Magia performs a two-step correction to the reference tissue mask (see below) before obtaining the input-function for modeling; the corrections makes the reference regions robust for many scanners and individuals. The MRI is also segmented into grey and white matter probability maps for spatial normalization ^15^. After modelling, the parametric images are spatially normalized and smoothed. In addition to the parametric images, Magia also calculates ROI-level parameter estimates for each study. Finally, the results are stored in a centralized archive in a standardized format along with visual quality control metrics, facilitating future population-level analyses.

**Figure 1.**
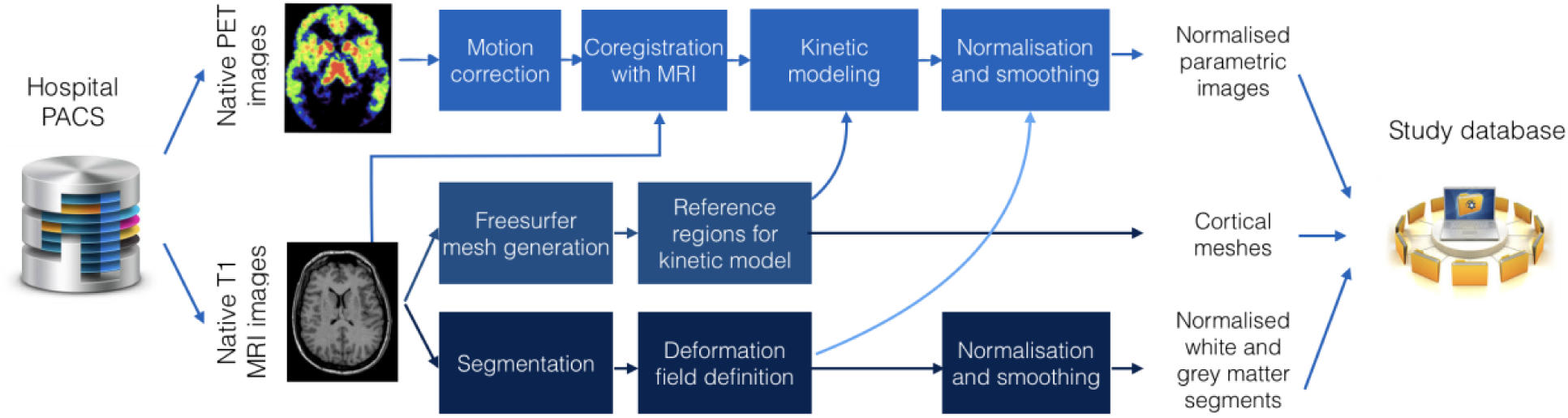
The Magia pipeline combining FreeSurfer cortical mesh generation and parcellation, T1 MRI image segmentation and normalization, automatic reference region and region of interest generation, and kinetic modeling.

The above-mentioned steps are only used when applicable. For example, for static images the frame alignment is skipped, and if there is no related MRI available, then a tracer-specific radioactivity template must be available to normalize the images. For all of the tracers included in this manuscript, such templates can be obtained from http://aivo.utu.fi/templates/. Magia also supports tracers that do not have a reference region. For such studies, the preprocessed plasma input must be available. Magia has default settings for preprocessing, modeling, and post-processing that have worked well during its development. However, Magia is also flexible in the sense that the user can override some of these options if needed.

### Validation data

To assess reliability of Magia we used historical control data using four radioligands with different targets and spatial distribution of binding sites: Dopamine D_2_R receptor antagonist [_11_C]raclopride, μ-opioid receptor agonist [_11_C]carfentanil, serotonin transporter ligand [_11_C]MADAM, and beta-amyloid ligand [_11_C]PIB. For each radioligand we selected 30 studies (Table 1). We generated reference regions for all the tracers using traditional manual methods and the new automatic method and compared the results.

**Table 1.** Summary of the studies. Scanners: HRRT (HRRT, Siemens Medical Solutions); PET/CT (Discovery 690 PET/CT, GE Healthcare); PET/MR (Ingenuity TF PET/MR, Philips Healthcare); GE Advance (GE Advance, GE Healthcare).

|  | [<sup>11</sup>C]carfentanil | [<sup>11</sup>C]raclopride | [<sup>11</sup>C]MADAM | [<sup>11</sup>C]PiB |
| --- | --- | --- | --- | --- |
| **N (female)** | 30 (12) | 30 (23) | 30 (17) | 30 (18) |
| **Age (mean, range)** | 32 (20 - 51) | 39 (20 - 60) | 42 (25 - 57) | 71 (66 - 80) |
| **Scanners** | HRRT | GE Advance | HRRT | HRRT |
|  | PET/CT | PET/CT |  |  |
|  | PET/MR | HRRT |  |  |
| **Data range (years)** | 2007 - 2016 | 1998 - 2014 | 2008 - 2015 | 2014 - 2016 |

### Manual reference region delineation

Five researchers with good knowledge of human neuroanatomy delineated reference regions for every study according to written and visual instructions (Figure 2a). Cerebellar cortex was used as a reference region for [_11_C]raclopride ^16^, [_11_C]MADAM ^17^ and [_11_C]PiB ^18^. For [_11_C]carfentanil, occipital cortex was used ^19^. The regions were drawn using CARIMAS (http://turkupetcentre.fi/carimas/) on three consecutive transaxial slices of T1-weighted MR images, which is the current standard manual method at Turku PET Centre. Cerebellar reference was drawn in cerebellar gray matter within a gray zone in the peripheral part of cerebellum, distal to the bright signal of white matter. The first cranial slice was placed below occipital cortex to avoid spill-in of radioactivity. Typically, this is a slice where the temporal lobe is clearly separated from the cerebellum by the petrosal part of the temporal bone. The most caudal slice was typically located in the most caudal part of the cerebellum. Laterally, venous sinuses were avoided to avoid spill-in during early phases of the scans. Posteriorly, there was about a 5 mm distance from cerebellar surface to avoid spill-out effects. Anteriorly, the border of the reference region was drawn approximately 2 mm distal to the border of cerebellar white and gray matter, except in the most caudal slice, where central white matter may no longer be visible.

**Figure 2.**
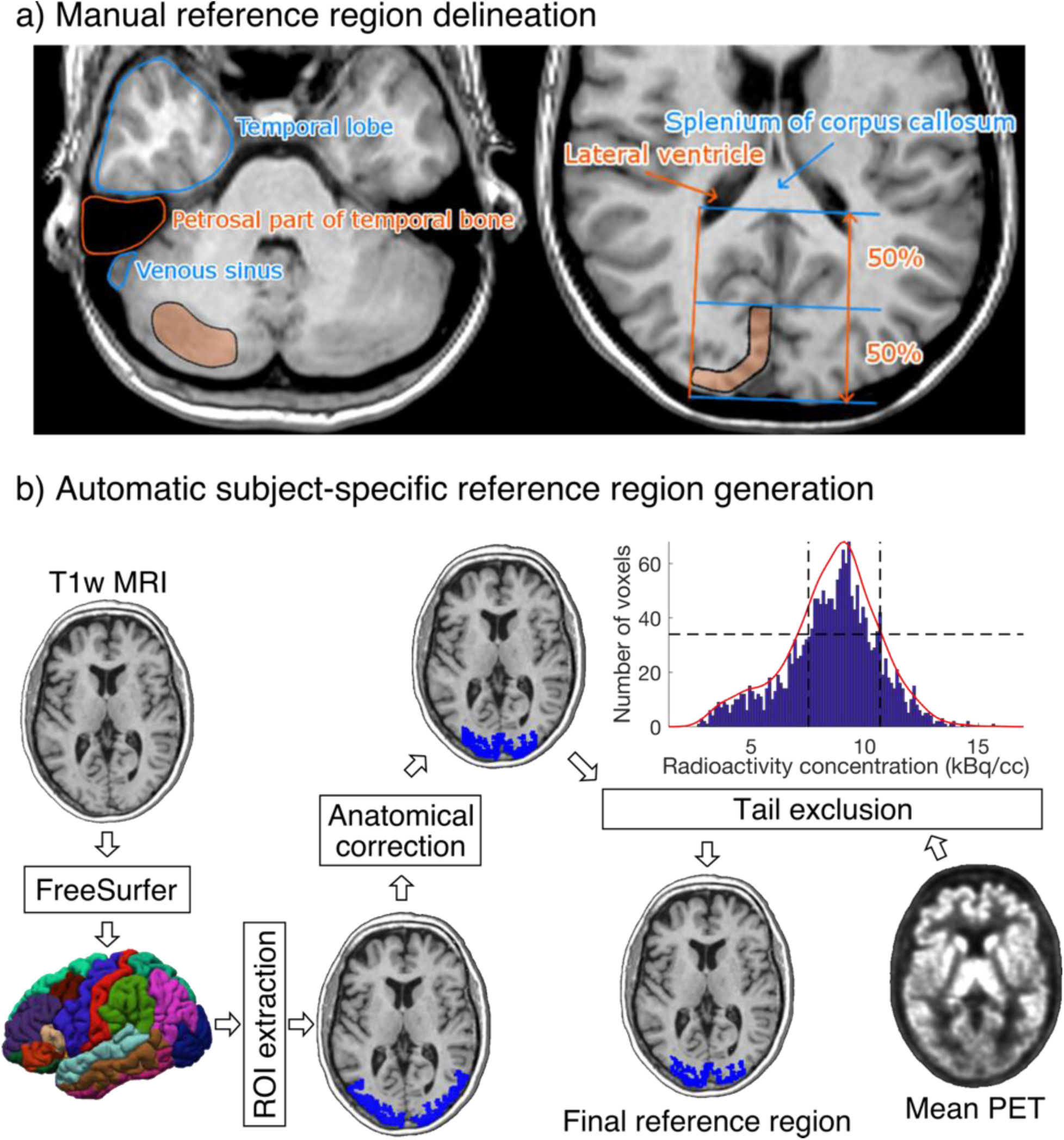
a) Visual instructions of the most cranial slice of manually delineated cerebellar (left) and occipital (right) reference regions. The reference regions were delineated on three consecutive transaxial T1-weighted MR images. Cerebellar reference region is shown on the left and occipital reference region on the right. b) The diagram shows how a T1-weighted MR image of an individual’s brain is processed to produce the final reference region. The shown example is from the [_11_C]carfentanil data set. The rectangles represent processing steps between inputs and outputs. The FreeSurfer step assigns an anatomical label to each voxel of the subject’s T1 weighted MR image. The ROI extraction step extracts a prespecified region of interest from FreeSurfer’s output. The anatomical correction removes voxels that are most likely to suffer from spillover effects; for [_11_C]carfentanil data this means voxels lateral to the lateral ventricles. In the tail-exclusion step, radioactivity distribution within the anatomically corrected reference region is estimated, and the voxels whose intensities are on the tail-ends of the distribution are excluded.

The occipital reference region was defined on three consecutive transaxial slices, of which the most caudal slice was the second-most caudal slice before cerebellum. The reference region was drawn J-shaped with medial and posterior parts. The reference region was drawn to roughly follow the shape of the cortical surface, but not individual gyri. The reference region was drawn approximately 1 cm wide with about 2 mm margin to the cortical surface to avoid spill-out effects. The anterior border of the reference region was placed approximately halfway between the posterior cortical surface and the splenium of corpus callosum. The posterolateral border of the reference region approximated the medial-most part of the posterior horn of the lateral ventricle.

### Automatic reference region generation

Figure 2b shows an overview of the automated reference-region-generation process. First, T1-weighted MR images were fed into FreeSurfer to provide subject-specific anatomical masks for cerebellar and occipital cortices. Second, an anatomical correction was applied to the FreeSurfer-generated reference region mask to remove voxels that, based on their anatomical location alone, are likely to suffer from spillover effects. For cerebellar cortex, the most important sources of spillover effects are occipital cortex and venous sinuses. Thus, the most outermost cerebellar voxels were excluded in the anatomical reference region correction. For occipital cortex, voxels that were lateral to the lateral ventricles were excluded. This is because the most lateral parts of the FreeSurfer-generated occipital cortex extend to areas with specific binding for [_11_C]carfentanil, and the lateral ventricles provide a reliable anatomical cut-off point for thresholding. Finally, the radioactivity concentration distribution within the anatomically corrected reference region was estimated, and the tails of the distribution were excluded. The lower and upper boundaries for the signal intensities were defined by calculating the full width at half maximum (FWHM) of the mean PET signal intensity distribution. This step ensures that the reference region will not contain voxels with atypically high or low radioactivity (e.g. signal from outside the brain). The automatic reference region generation process thus combines information from anatomical brain scans and the PET images to get a reliable estimate of nonspecific binding.

### Quantifying operator-dependent variability

We first investigated how subjective choices in manual reference-region delineation translate into differences in reference region masks, reference-region time-activity curves, and outcome measures. Anatomical differences in reference region masks were assessed in two ways: First, we calculated within-study spatial overlap between the manual reference regions. The spatial overlap was calculated in two stages: it was first calculated separately for all different manual reference region pairs, and those numbers were then averaged over to obtain a summary statistic for each study. Second, we investigated the differences in volumes of the manually delineated reference regions using intra-class correlation coefficient (ICC). To estimate ICC, we first estimated a random effects model y ∼ 1 + (1 | z), where y is the variable of interest, and z is the grouping variable (here either study or operator), and then calculated the proportion of total variance explained by the variance of the random effect -component ^20^. Calculated this way, ICC is restricted to between 0 and 1. The R package brms (https://cran.r-project.org/package=brms) was used to estimate the models, and the R package performance (https://easystats.github.io/performance/index.html) was used to estimate ICC.

Differences in reference region TACs were assessed by calculating area under the curve (AUC) of them. Prior to the ICC analysis, we standardized all the AUCs with the mean radioactivity within union of all manually delineated reference regions. This standardization removes uninteresting between-study variance resulting from different scanners, body masses and injected doses. The operator-caused variation in outcome measures was also assessed using ICC.

### Volumetric similarity of the manual and automatic reference regions

We compared the volumes of reference regions to assess whether the two techniques generate reference regions of systematically different sizes. For each study, we calculated the mean volume from all manually delineated reference regions and compared it to volume of the Magia-derived reference region. We also quantified the anatomical overlap between the manually and the automatically derived reference regions. The overlap was defined as ratio between the number of common voxels and the number of manual voxels. For each study, the overlap was first calculated separately for every manually delineated reference region after which the mean overlap was calculated.

### Similarity of the reference region radioactivity concentrations

A functionally homogenous region should have approximately Gaussian distribution of radioactivity measured with PET ^21^. Functional homogeneousness was assessed using radioactivity distributions within the reference regions. The automatically and manually derived reference region masks were used to extract radioactivity concentration distribution within the reference regions. The study-specific manual distributions were averaged over the manual drawers to provide a single manual distribution for each study. The radioactivity concentrations were converted into SUV, after which the distributions were averaged over studies to provide tracer-specific distributions. Mean, standard deviations, mode, and skewness of the distributions were used to quantify the differences in the distributions.

### Similarity of the reference region time-activity curves

We compared the similarity of the automatically and manually delineated reference region time-activity curves (TACs). For each study, the manual reference region TAC was defined as the average across the manual TACs to minimize the subjective bias in adhering to the instructions for manual reference region delineation. Activities were expressed as standardized uptake values (SUV, g/ml) which were obtained by normalizing tissue radioactivity concentration (kBq/ml) by total injected dose (MBq) and body mass (kg), thus making the different images more comparable to each other. To assess the similarity of the shapes of reference region TACs, we calculated Pearson correlations between the manually and automatically delineated TACs for each tracer. Bias was assessed using area under curve (AUC).

### Assessing similarity of the outcome measures

We used nondisplaceable binding potential (*BP_ND_*) to quantify uptakes of [_11_C]carfentanil, [_11_C]raclopride and [_11_C]MADAM. It reflects the ratio between specific and nondisplaceable binding in the brain. The binding potentials were calculated using simplified reference tissue model whose use has been validated for these tracers ^16, 17, 19^. SUV-ratio between 60 and 90 minutes was used to quantify [_11_C]PiB uptake ^18^. All the studies were first processed using Magia. To obtain the outcome measures resulting from manually delineated reference regions the procedure was repeated with the only exception of replacing the automatically generated reference regions with a manually generated reference region. Thus, the only differences observed in the uptake estimates originate from differences in the reference regions. We estimated the outcome measures in one representative region of interest (ROI) for each tracer, and also calculated parametric images. The ROIs were extracted from the FreeSurfer parcellations.

## Results

### Operator-dependent variation

The influence of different operators on reference regions, reference region time-activity curve, and outcome measures are presented for each tracer (Table 2). The spatial overlap between the manually delineated masks was modest, as the maximum overlap was 41 % for [_11_C]raclopride studies, while the overlap for the other tracers was 14 to 22 %. The reference region volumes were most similar between the operators for [_11_C]carfentanil (ICC = 69 %). For [_11_C]PiB and [_11_C]MADAM, the reference region volume ICC was at most 5 %, while the operator had up to 62 % influence on the reference region’s volume for [_11_C]PiB. The reference region TAC AUCs varied substantially especially for [_11_C]carfentanil, while for other tracers the AUCs were highly similar across operators. The operator-caused variation was less substantial in the outcome measures, but still significant particularly for [_11_C]carfentanil.

**Table 2.** Operator-caused variation in basic characteristic derived from the reference region masks.

| Tracer | Spatial overlap (%) | Variance explained by study (%) |  |  | Variance explained by operator (%) |  |  |
| --- | --- | --- | --- | --- | --- | --- | --- |
|  |  | Reference region volume | Ref-TAC AUC | Outcome measure | Reference region volume | Ref-TAC AUC | Outcome measure |
| [<sup>11</sup>C]carfentanil | 22 | 69 | 33 | 47 | 9 | 22 | 21 |
| [<sup>11</sup>C]raclopride | 41 | 43 | 77 | 96 | 28 | 2 | 1 |
| [<sup>11</sup>C]PiB | 14 | 2 | 94 | 95 | 62 | 1 | 1 |
| [<sup>11</sup>C]MADAM | 18 | 5 | 54 | 65 | 37 | 3 | 7 |

### Differences between manually and automatically produced reference regions

#### Differences in reference region masks

We first compared the anatomical similarities between the automatically and manually delineated reference regions. For each tracer, automatic reference regions were consistently larger than manually derived reference regions (Figure 3 and Figure 4a). In four [_11_C]carfentanil studies at least one of the manually drawn reference region was larger than the automatic occipital reference region. Magia-generated cerebellar reference regions were always larger than mean manual cerebellar reference regions. The automatically produced reference regions are naturally larger than the manually delineated ones because manual delineation requires mechanic work from highly trained individuals, thus providing a cost to the size of the regions.

**Figure 3.**
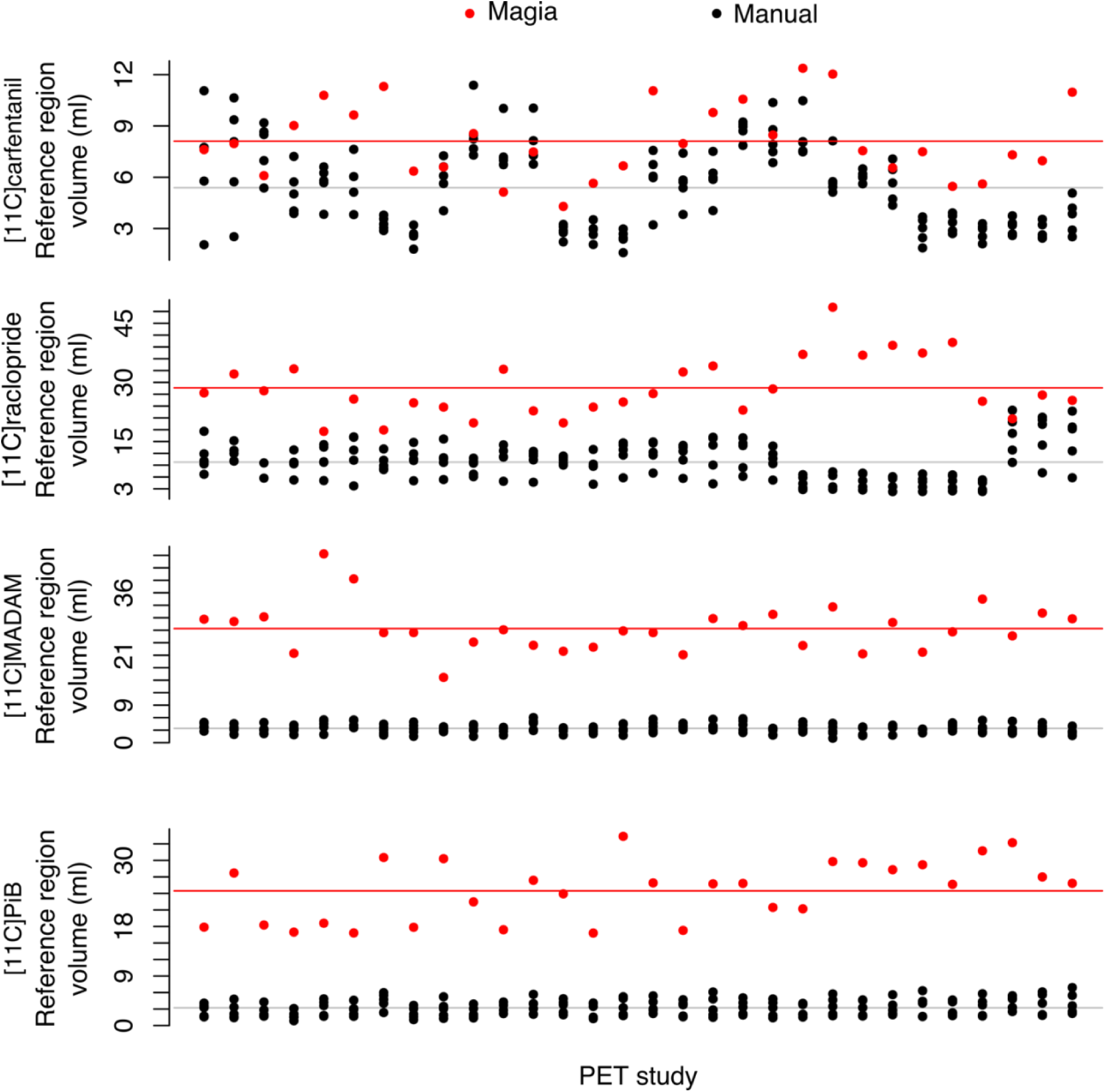
Differences in volumes of manually and automatically produced reference regions.

**Figure 4.**
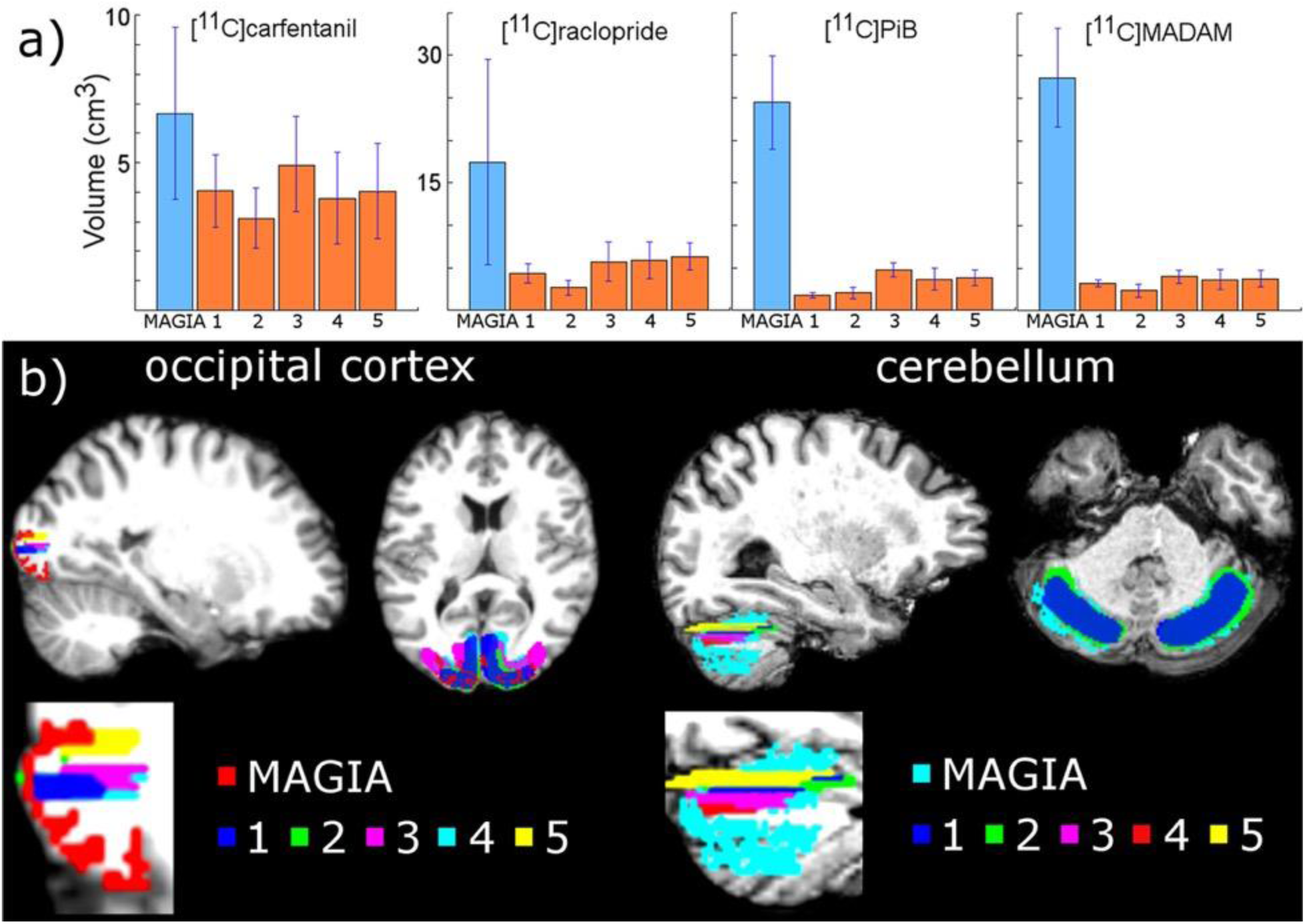
a) Mean volumes of Magia-generated reference regions compared to mean volumes of manually delineated reference regions. b) Visual example of Magia-generated and manual reference regions for one study.

Next, we determined whether the Magia-derived reference regions overlap with the manually drawn reference regions. Automatic occipital reference region for [_11_C]carfentanil overlapped only 14 % with manual occipital reference region. However, automatic cerebellar reference regions overlapped with manual reference regions by 55 %, 59 % and 61 % for [_11_C]raclopride, [_11_C]MADAM and [_11_C]PiB, respectively.

#### Differences in reference region SUV distributions

The overlap between the manual and automatic radioactivity distributions was approximately 90 % for all tracers (Figure 5). All distributions were unimodal and highly symmetric for all tracers. The means of the distributions were practically equal (maximum difference of 0.07 %). The standard deviations of the distributions differed by 14 %, 11 %, 12 % and 18% for [_11_C]carfentanil, [_11_C]MADAM, [_11_C]PIB and [_11_C]raclopride, respectively. The modes of the automatically and manually derived distributions were 1.5 and 1.55 for [_11_C]carfentanil, 1.95 and 2.05 for [_11_C]MADAM, 1.65 and 1.70 for [_11_C]PIB, and 1.35 and 1.35 for [_11_C]raclopride. Thus, the maximum difference was less than 5 %. The skewnesses of the Magia-derived and manually derived distributions were 1.2 and 0.9 for [_11_C]carfentanil, 1.3 and 1.2 for [_11_C]MADAM, 2.0 and 1.6 for [_11_C]PIB, and 2.4 and 2.0 for [_11_C]raclopride.

**Figure 5.**
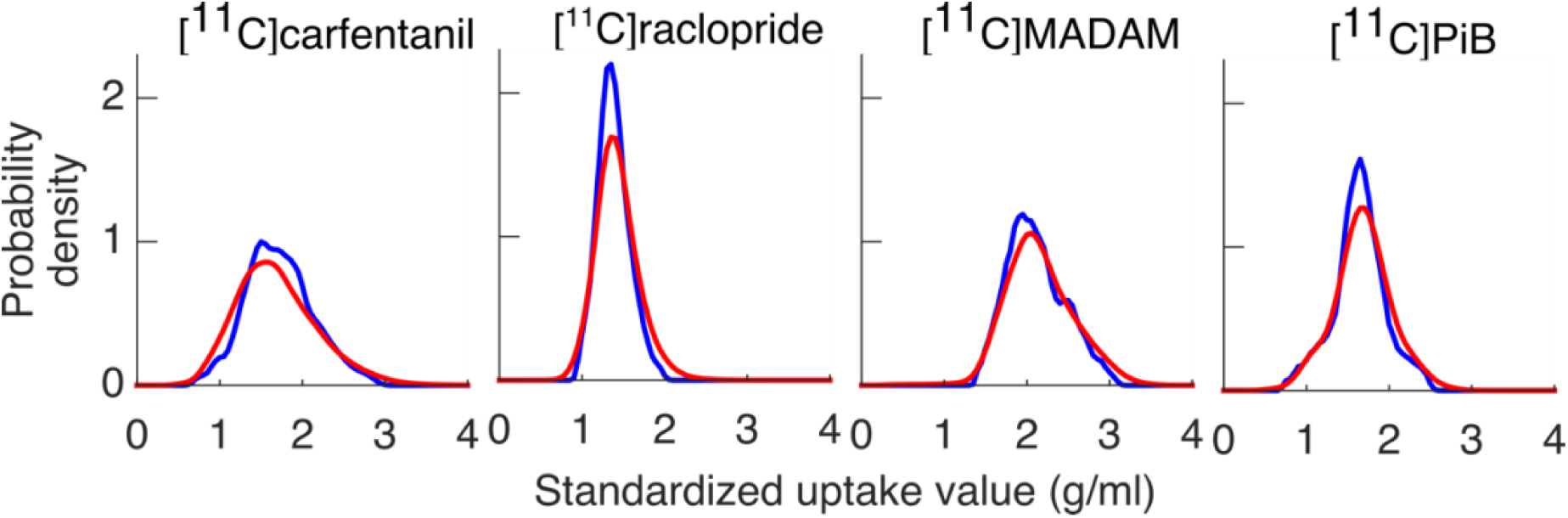
Between-subject average reference region radioactivity distributions. Blue = Magia; red = manual.

### Differences in reference region time-activity curves

The Magia-produced time-activity curves were on average very similar to the average TACs calculated based on the manually delineated reference regions (Figure 6). The Pearson correlation coefficients were above 0.99 for all tracers. Figure 7 shows how the Magia-derived reference region time-activity curve AUCs compare against the manually obtained results. For [_11_C]carfentanil, the between-study AUC means were practically identical (< 1 %). The Magia-produced reference regions had 2.6 %, 1.1 %, and 1.8 % lower AUCs than the manual reference regions for [_11_C]raclopride, [_11_C]MADAM, and [_11_C]PiB, respectively.

**Figure 6.**
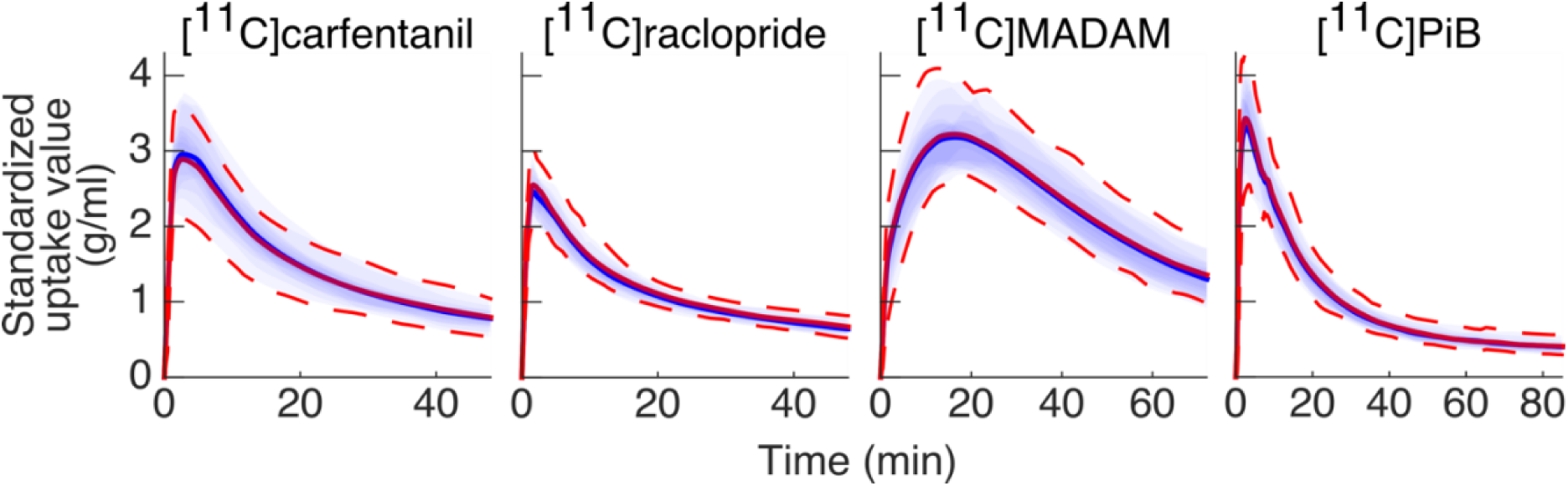
Between-subject mean time-activity curves. Blue = Magia; red = manual.

**Figure 7.**
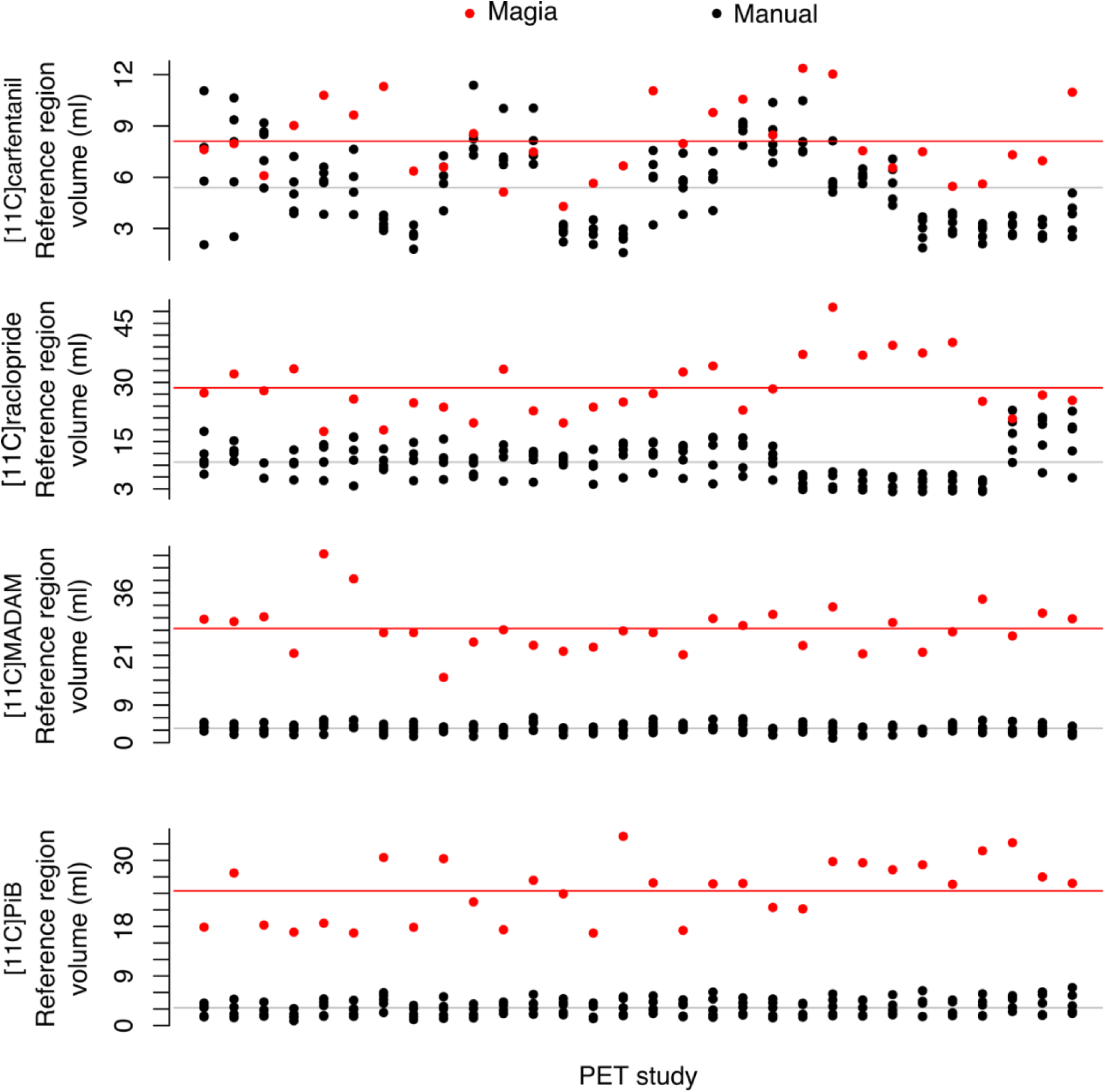
Comparison of reference region volumes.

### Differences in outcome measures

Pearson correlation coefficients between the mean of manual outcome measures and the Magia-derived outcome measures were 0.79, 0.98, 0.84, and 0.99 for [_11_C]carfentanil, [_11_C]raclopride, [_11_C]MADAM, and [_11_C]PiB, respectively. The outcome measures derived using automatic and manual methods are visualized in Figure 9 in one representative region of interest, and the relative bias in the whole brain between them is visualized in Figure 8b. For [_11_C]carfentanil and [_11_C]PiB Magia produced basically no bias (less than 1 %). For [_11_C]MADAM, Magia produced up to 3–5 % higher binding potential estimates in regions with high specific binding. In cortical regions with low specific binding, the bias was over 10 %. For [_11_C]raclopride, Magia produced approximately 4–5 % higher binding potential estimates in striatum. In thalamus, the bias was 8–10 %. Elsewhere in the brain the bias varied considerably between 13–20 %. These differences were all statistically significant (FWE-corrected voxels, p < 0.05). For both [_11_C]MADAM and [_11_C]raclopride, the relative bias decreased significantly with increasing binding potential (Figure 8c).

**Figure 8.**
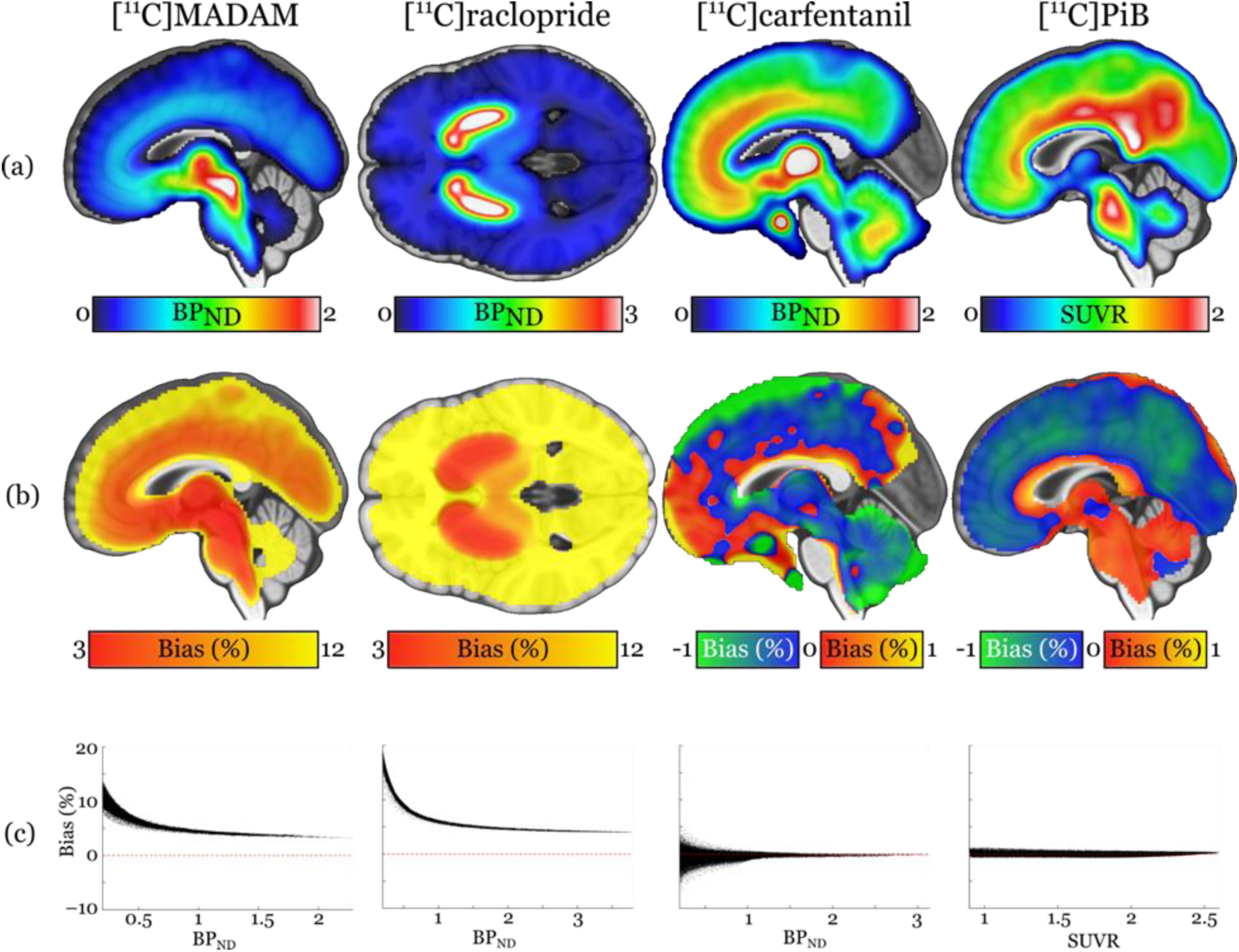
(a) Visualization of the outcome measure distributions for each tracer. (b) Maps visualizing the relative biases of the Magia-derived outcome measures compared to the averages obtained by manual reference region delineation. The manual method is here presented as the ground truth, because the manual outcome for each scan is an average over five individual estimates, while the Magia result relies on a single estimate. (c) Associations between the outcome measure magnitude and relative bias.

**Figure 9.**
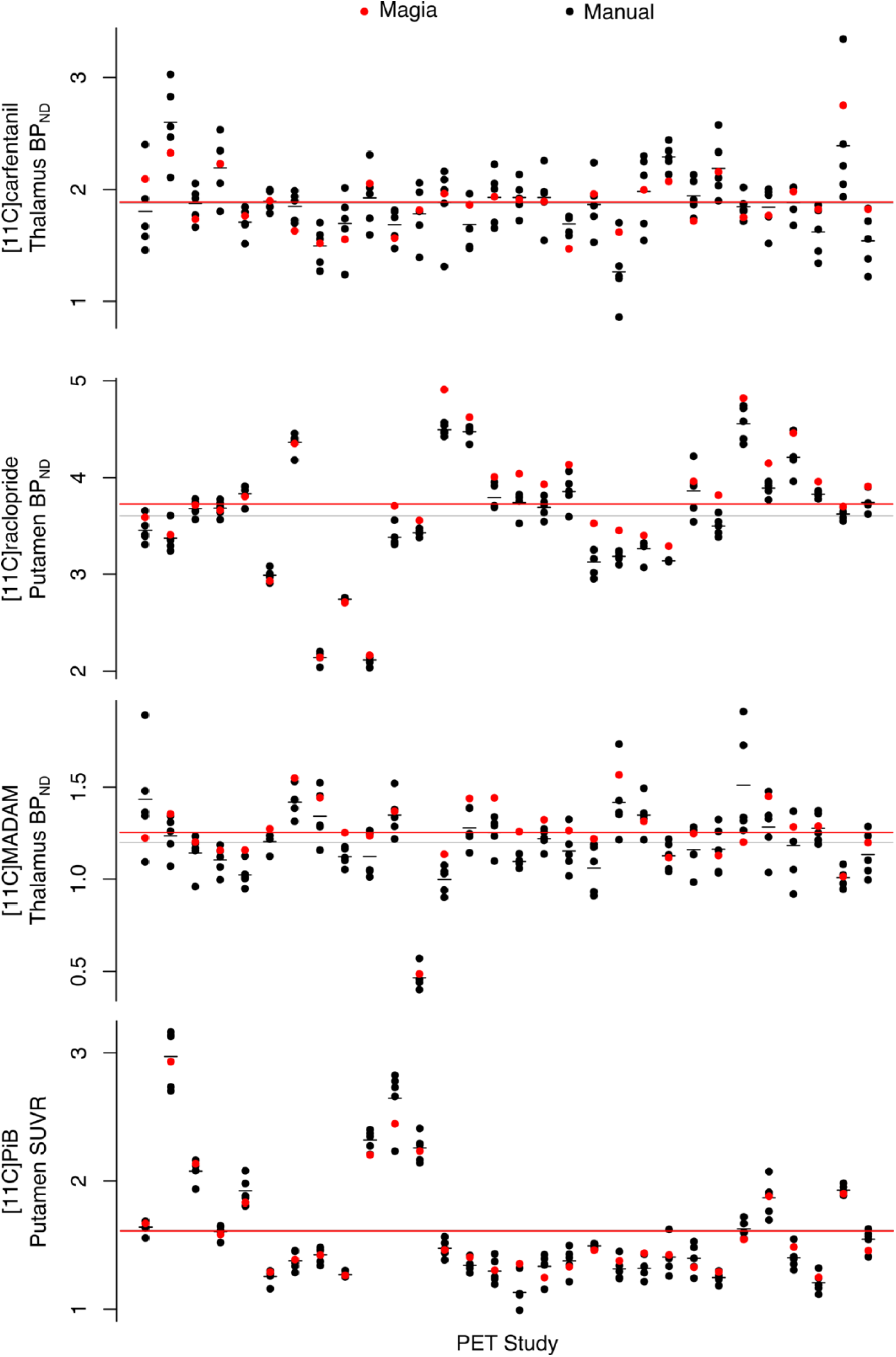
Comparison of Magia-derived outcome measures against manually obtained ones.

## Discussion

We established that the automated Magia pipeline produces consistent estimates of radiotracer uptake for all the tested ligands, with very little or even no bias in the outcome measures. As expected, the manual delineation method suffered from significant operator-dependent variability, highlighting the importance of standardization of the process. The consistency coupled with significant gains in processing speed suggests that Magia is well suited for automated analysis of brain-PET data for large-scale neuroimaging projects.

### Outcome measures can substantially depend on who delineated the reference region

We estimated the amount of operator-dependent variation in outcome measures. Despite all operators drawing the ROIs using the same instructions (presented both verbally and as visual / written instructions available for reference while working) the ICC analyses show that for [_11_C]carfentanil, the variation produced by different operators is significant. Approximately half of the variation between studies (47 % vs. 21 % of total variance explained) was due to operator variability. These numbers were worst for [_11_C]carfentanil, indicating that for [11C]carfentanil PET studies the subjective variation in manual ROI delineation (e.g. which transaxial slices to use, how to define ROI boundaries etc.) significantly influence the magnitude of binding potential estimates. Out of the tracers using cerebellar cortex as the reference region, [_11_C]MADAM had the lowest ICC with 65 %. For [_11_C]raclopride and [11C]PiB the ICCs were over 95 % and the operator’s effect was less than 2 %, indicating that for these tracers manual delineation of reference regions may not be as crucial source of variation.

These differences between tracers likely reflect differences in uniformity of the PET signal within the reference region. If the reference region were perfectly homogenous with respect to the PET signal, it would not matter at all which voxels to choose. In reality, however, the PET signal is highly heterogenous. For example, the PET signal depends on the transaxial slices used. Presumably these heterogeneities are substantial for [_11_C]carfentanil and, to a lesser extend, [_11_C]MADAM, while the PET signal from cerebellar cortex using [_11_C]raclopride and [_11_C]PiB is significantly more homogenous. Indeed, the spatial overlap between the manually delineated reference region was higher for [_11_C]carfentanil (22 %) than for [11C]PiB (14 %), suggesting that even small differences in spatial overlap translate into substantial differences in binding potential for [_11_C]carfentanil.

While the between-operator variance in outcome measures was substantial, the influence of the operator on reference TAC AUCs was even larger. For all the tracers, the ICC of outcome measures was higher than the ICC for reference TAC AUCs. For example, while [_11_C]raclopride *BP_ND_* was barely influenced by the individual manually delineating the reference region, the ICC for [_11_C]raclopride reference TAC AUC was only 77 %, almost 20 %-units less than for *BP_ND_*. Thus, even the reference region TACs for [_11_C]raclopride were not remarkably consistent between the operators, further highlighting the sensitivity of the delineation process. The size of the reference region was not even consistent for each individual performing the delineation. The highest proportion of variance explained by individual operators was observed for [_11_C]PiB with 62 %. For [_11_C]carfentanil, on the other hand, the operator only explained 21 % of the variation in reference region volumes, showing that even single individuals can define reference regions that remarkably vary in their size, despite detailed written and visual instructions. This result highlights the need for reference-region generation processes that do not suffer from subjectivity.

### Reliability of Magia’s uptake estimates

Importantly, Magia produced parameter estimates consistent with the *averaged* manual estimates (Pearson correlation coefficients > 0.78 for all tracers). This suggests that i) even though individual operators yield different output metrics these are sampled from the same true parameter space, which ii) is in turn accurately reflected by the Magia output. There was no systematic bias for [_11_C]PiB SUVR and [_11_C]carfentanil *BP_ND_*. For [_11_C]PiB, the difference between the manual and automatic SUVR estimates fluctuated randomly around zero. Because SUVR was used to quantify [_11_C]PiB uptake, the random fluctuation was independent of brain region. For [_11_C]carfentanil, the random fluctuation was slightly greater in low-binding regions (but still within +/-5 %). In contrast to [_11_C]PiB and [_11_C]carfentanil, there were systematic differences between the manual and automatic binding potential estimates for [_11_C]raclopride and [_11_C]MADAM. For both tracers the bias decreased as a function of specific binding, and in high-binding regions (*BP_ND_* > 1.5) the bias was less than 5 %. Even if the bias increased sharply with decreasing binding potential, the problematic regions are not typically considered very interesting because of their poor signal-to-noise ratio.

The systematic bias for [_11_C]MADAM and [_11_C]raclopride is also reflected in the small differences in reference tissue TACs. For the tracers using cerebellar reference region, Magia-derived reference tissue TACs had 2-3 % lower AUCs. The peaks of the TACs were also slightly lower. For [_11_C]PiB, the bias did not propagate into outcome measures because the SUV-ratio was calculated between 60 and 90 minutes when there was no bias in TACs. Because binding potential reflects the ratio between specific binding and unspecific binding (obtained from reference tissue), the reference TAC AUCs directly propagate into biases in binding potentials. Thus, these data indicate that Magia may produce slightly higher binding potential estimates than traditional methods at least if cerebellar cortex is used as the reference region. These data do not however imply that the bias should be regarded as error: In fact, Magia produces significantly larger reference regions, and consequently the reference tissue TACs are less noisy. This is desirable, because the noise in input function influences model fitting. However, the bias also means that Magia-produced estimates should not be combined with estimates produced with other methods.

### Functional homogeneity of the reference regions

We tested whether the assumption of homogenous binding within the reference regions holds for both automatic and manual reference regions. A homogenous source region should produce unimodal and approximately symmetric radioactivity distributions ^21^. Between-study average distributions were unimodal and symmetric for all tracers for both the manual and automatic method. The distribution means were practically identical, but the modes were 1–2 % higher for Magia. The manual distributions were slightly wider (the standard deviations were approximately 15 % larger) because Magia cuts the distribution tails. The manual distributions were also slightly less skewed. Because averaging distributions tends to make them more Gaussian, this difference probably arises from the fact that the manual distributions that were used in the comparison were defined as an average over the five distributions delineated by the independent operators. The distribution overlaps were approximately 90 % for all tracers. In sum, these results show that the Magia-generated reference region radioactivity distributions satisfy the requirement of functional uniformity.

### Reference tissue time-activity curves

Despite their topographical differences, the automatically and manually produced reference regions yielded very similar time-activity curves. For all tracers, the Pearson correlation coefficient between average automatic and manual reference tissue TACs was above 0.99. The TAC shapes were thus in excellent agreement. For [_11_C]carfentanil, also the AUC of reference region TACs were highly similar. The AUCs of cerebellar time-activity curves were 2-3 % lower for Magia, indicating that the cerebellar automatic TACs were slightly negatively biased compared to their manual counterparts. These data do not directly indicate which method produced more realistic time-activity curves. However, because the Magia-generated cerebellar reference regions were without exception substantially larger than their manual counterparts, the time-activity curves of Magia presumably have higher signal-to-noise ratio, suggesting that the Magia-derived metrics may compare favorably against the manually obtained metrics.

### Solving time constraints in processing of PET data

On average, drawing the reference region for a single subject took around fifteen minutes, and without any automatization the modeling and spatial processing of the images standard tools (e.g. PMOD or Turku PET Centre modelling software) takes on average 45 minutes. In contrast, it takes less than five minutes to set Magia running for a single study. Although the time advantage– roughly an hour per study–gained from automatization is still modest in small-scale studies (e.g. three eight-hour working days for a study with 24 subjects) the effect scales up quickly, and manual modeling of a database of just 400 studies would take already fifty days. This is a significant investment of human resources, in particular if the analyses have to be redone later with, for example, different modeling parameters requiring repeating of at least some parts of the process.

### Standardized processing creates novel opportunities for data sharing

Functional neuroimaging community has already established standardized pipelines for preprocessing fMRI data. However, a publicly available pipeline that automatically produces the outcome measures from PET images in a standardized fashion has been lacking. Of course, also the brain PET community has used standardized methods as much as possible. Magia takes the standardization to extreme by providing a fully automated and standardized analysis option for brain PET studies, especially for studies where reference-region based modeling can be used. The increased standardization decreases variance resulting from subjective choices in the analysis process, thus reducing unnecessary variation also in population level analyses and subsequently increasing statistical power. Importantly, standardized processing provides novel opportunities for data sharing between different sites, because it is easier to combine data from different sites if the data have been processed using exactly the same methods.

### Limitations

Magia is currently fully automatic only for tracers for which a reference region exists. However, even for blood-based inputs, Magia requires minimal user intervention, as Magia can read in the input from the appropriate location. Magia was originally developed with the assumption that a T1-weighted MR image is available for each subject (for reference region delineation and spatial normalization). Because this assumption limited the applicability of the approach for reanalysis of some historical data, Magia can now also use templates for ROI definition and tracer-specific radioactivity templates for spatial normalization. Templates for each of the tracers used in this manuscript are available in https://github.com/tkkarjal/magia/tree/master/templates, and Magia can use whatever templates the user may have available. Thus, availability of MRI is not necessary, but it is strongly recommended because most of the testing has been done with MRI-based processing, and because the ROIs as well as reference regions can then generated in the native space. Finally, Magia processes the studies independently of each other. Within-subject designs would benefit from consideration of multiple images per participant, but this is currently not possible.

## Conclusions

Magia is a standardized and fully automatic analysis pipeline for processing brain PET data. By standardizing the reference region generation process, Magia eliminates operator-dependency in producing outcome-measures. For [_11_C]carfentanil that uses occipital cortex as the reference region, the reduced variance comes with no cost for bias in *BP_ND_*. The SUVR estimates were also unbiased for [_11_C]PiB. [_11_C]raclopride and [_11_C]MADAM *BP_ND_* were slightly overestimated. However, compared to the variance resulting from operator dependency, this bias was negligible and may actually favor Magia. In any case, bias is meaningless in most population-level analyses. Magia enables standardized analysis of brain PET data, facilitating shift towards larger samples and more convenient data sharing across research cites.

## Acknowledgements

This work was supported by the Academy of Finland grants #265915 and #294897 to LN, and Sigrid Juselius Foundation separate grants to LN and JR, Päivikki and Sakari Sohlberg Foundation to ToK, Finnish Cultural Foundation Varsinais-Suomi Regional Fund to ToK, State Research Funding (VTR) for Turku University Hospital to ToK, State Research Funding (VTR) for Turku University Hospital to JR

## Notes

#### Summary of Updates

Added analyses regarding operator-dependency

https://github.com/tkkarjal/magia

